# miR-203 imposes an intrinsic barrier during cellular reprogramming by targeting NFATC2

**DOI:** 10.1101/2020.06.02.131136

**Authors:** María Salazar-Roa, Sara Martínez-Martínez, Osvaldo Graña-Castro, Mónica Álvarez-Fernández, Marianna Trakala, Juan-Miguel Redondo, Marcos Malumbres

**Affiliations:** Cell Division and Cancer Group, Spanish National Cancer Research Centre (CNIO), Madrid, Spain; Genetic Regulation, Vascular Remodeling and Inflammation group, Spanish National Cardiovascular Research Center (CNIC), Madrid, Spain; Bioinformatics Unit, CNIO, Madrid, Spain

**Author notes:** Correspondence: M.M., Centro Nacional de Investigaciones Oncológicas, Melchor Fernández Almagro 3, E-28029, Madrid.

## Abstract

Cellular reprogramming from somatic to pluripotent cells is the basis for multiple applications aimed to replace damaged tissues in regenerative medicine. However, this process is limited by intrinsic barriers that are induced in response to reprogramming factors. In this manuscript we report that miR-203, a microRNA with multiple functions in differentiation and tumor suppression, acts as an endogenous barrier to reprogramming. Genetic ablation of miR-203 results in enhanced reprogramming whereas its expression prevents the formation of pluripotent cells both in vitro and in vivo. Mechanistically, this effect correlates with the direct repression of NFATC2, a transcription factor involved in the early phases of reprogramming. Inhibition of NFATC2 mimics miR-203 effects whereas NFATC2 overexpression rescues inducible cell pluripotency in miR-203-overexpressing cultures. These data suggest that miR-203 repression may favor the efficiency of reprogramming in a variety of cellular models.

## Introduction

Cellular reprogramming is executed by complex gene regulatory networks ultimately leading to stable epigenetic modifications that determine a pluripotent cell state (Schmidt & Plath, 2012, Takahashi, 2014, Takahashi & Yamanaka, 2006). The induction of cellular reprogramming for specific transcription factors typically results in stress responses that limit the efficacy of this process, including senescence, p53- and CDKN2-pathways as well as epigenetic barriers (Banito, Rashid et al., 2009, Gascon, Masserdotti et al., 2017, Haridhasapavalan, Raina et al., 2020, Kawamura, Suzuki et al., 2009, Marion, Strati et al., 2009, Utikal, Polo et al., 2009, Zhuang, Li et al., 2018). Despite the advances in the last decade, reprogramming from somatic cells is still an inefficient process, limiting the application of this technology in regenerative medicine.

Several microRNAs have emerged as critical regulators of cellular reprogramming by either favoring or limiting this process from different cell types (Ishtiaq, Waseem et al., 2018, Srivastava & DeWitt, 2016). miR-203 was originally identified as a microRNA highly induced during differentiation from skin progenitors by targeting the transcription factor p63 (Jackson, Zhang et al., 2013, Lena, Shalom-Feuerstein et al., 2008, Yi, Poy et al., 2008). This microRNA was also found to be inactivated in tumors by genetic and epigenetic mechanisms and it has been proposed to be involved in protecting from tumor development and metastasis through the repression of multiple targets (Bueno, Perez de Castro et al., 2008, Michel & Malumbres, 2013). More recently, we reported the surprising observation that transient exposure of already-established induced pluripotent stem cells (iPSC) or embryonic stem cells (ESC) to miR-203 enhances the ability of these pluripotent cells to differentiate into multiple cell lineages and to reach further maturation properties (Salazar-Roa, Trakala et al., 2019). However, the relevance of this microRNA during reprogramming has not been tested. In this manuscript, we describe a dramatic effect of miR-203 in preventing reprogramming of somatic cells both in vitro and in vivo. miR-203 overexpression during reprogramming from primary fibroblasts results in reduced levels of the transcription factor NFATC2, a protein known to be required for early steps during reprogramming. On the other hand, genetic ablation of miR-203 results in enhanced reprogramming in vitro in the presence of high NFATC2 protein levels, suggesting that miR-203 functions as an endogenous barrier to cellular reprogramming.

## Results and Discussion

### miR-203 prevents cellular reprogramming from fibroblasts

We first generated a conditional knockout mouse model in which the miR-203 encoding gene can be eliminated after expression of Cre recombinase (Figure EV1A). We also made use of an inducible knock-in model in which miR-203 can be ubiquitously induced from the *Cola1* locus upon doxycycline treatment [*Cola1*(tet-miR-203) allele (Salazar-Roa et al., 2019)]. To evaluate the role of miR-203 on cellular reprogramming, we transduced wild-type, *miR-203*(−/−) or *Cola1*(tet-miR-203/tet-miR-203); *Rosa26*(rtTA/rtTA) [referred to as *Cola1*(miR-203) from now on] mouse embryonic fibroblasts (MEFs) with lentiviral vectors expressing Oct4, Sox2, Klf4 and cMyc (FUW-OSKM, non-inducible version)(Takahashi & Yamanaka, 2006). For inducing transient miR-203 over-expression in *Cola1*(miR-203) MEFs, cell cultures were treated with Doxycycline (1 mg/ml) during the complete reprogramming process. Doxycycline was added also to miR-203 wild-type and knock-out MEF cultures, as a control. Surprisingly, we found that the reprogramming process was notably accelerated in *miR-203*(−/−) cells, whereas miR-203 over-expression was detrimental for reprogramming (Figure 1A). Analysis of the reprogramming process by RNA sequencing revealed a clear evolving transcriptomic pathway from MEFs at day 0 to the final iPS cells (Figure 1B). Interestingly, whereas the three different lines [*miR-203*(+/+), *miR-203*(−/−), and *Cola1*(miR-203)] clustered together at day 0, they started to diverge at later time points, with miR-203-overexpressing cells remaining apart from full-developed iPSCs and *miR-203*(−/−) cells moving closer to full-developed iPSCs at early time-points. These differences were obvious when a previously-defined pluripotency-associated signature (Chung, Lin et al., 2012) was tested in these cultures (Figure 1C). In agreement with these observations, some critical genes involved in reprogramming, such as *Nanog, Dppa5 (Esg1), Eras, Khdc3, Fgf4* and *Gdf3*, were prematurely induced in *miR-203*(−/−) cultures, while miR-203 over-expression prevented their induction both at the transcript and protein levels (Figure 1D, E and Figure EV1B). Gene ontology analysis of genes significantly down-regulated in miR-203 over-expressing cells and significantly up-regulated in miR-203 knock-out cells suggested similar activation of cellular processes related to development and stem cell maintenance (Figure EV1B, C and Appendix Tables S1,2). Altogether, these data suggest a critical role for miR-203 in preventing reprogramming by altering the transcriptional changes necessary for the progression from somatic to reprogrammed cell fate, at least in fibroblasts.

**Figure 1.**
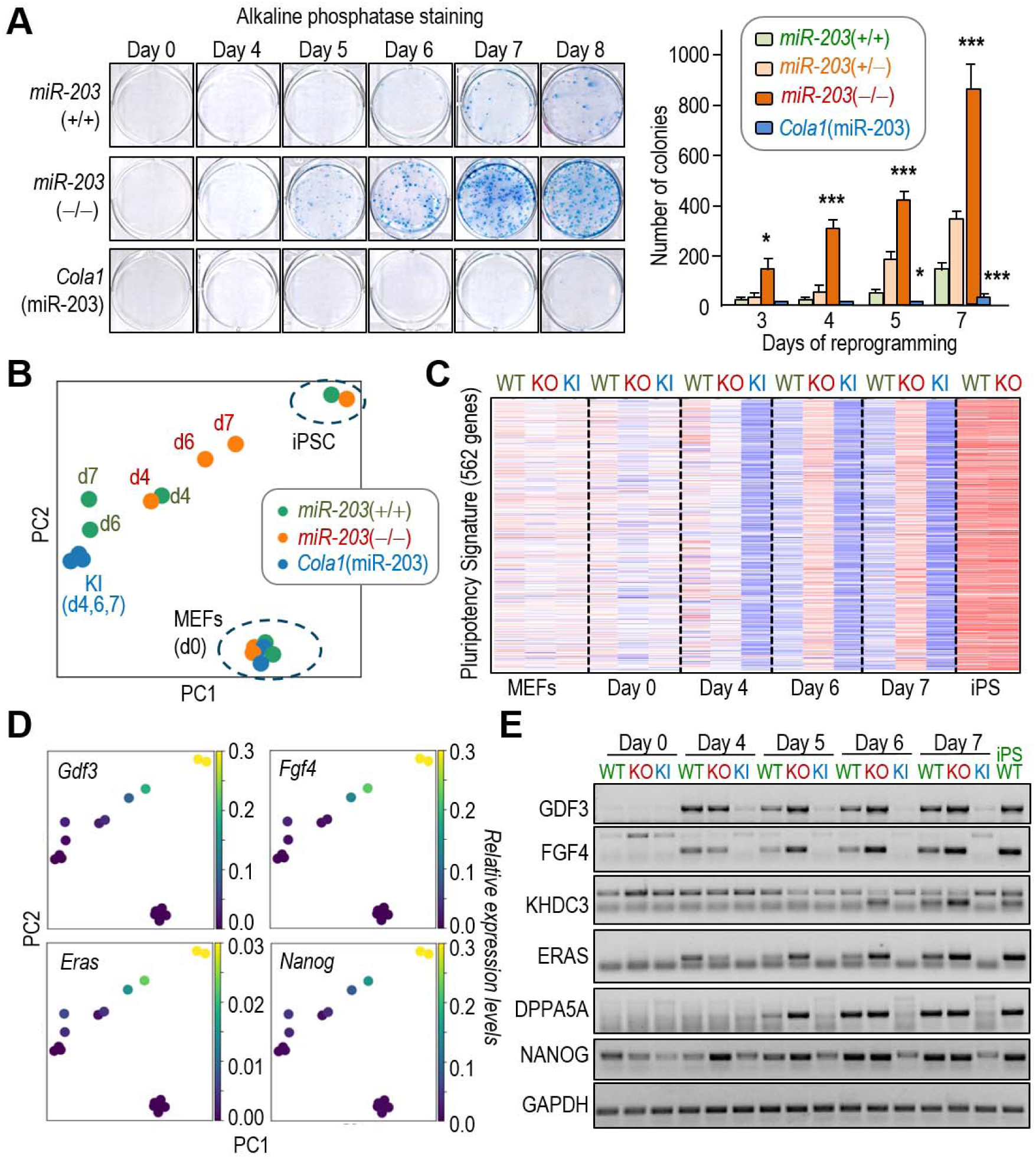
Efficient reprogramming in miR-203-deficient cells. **A**, MEFs of the indicated mouse lines were transduced with non-inducible Oct4-Sox2-Klf4-cMyc (OSKM) lentiviral vectors. Doxycycline was added during the complete reprogramming process to the three groups, leading to miR-203 over-expression in the *Cola1*(miR-203/miR-203); *Rosa26*(rtTA/rtTA) MEFs [referred to as *Cola1*(miR-203)]. The quantification of the number of alkaline phosphatase (AP)-positive colonies is shown to the right. Values correspond to the average and s.d. (n= 5 independent MEF isolates per line). *, *P*<0.05; **, *P*<0.01; ***, *P*<0.001 (Student’s t-test; unpaired; two-tailed). **B**, Principal Component Analysis of RNAseq data from all the samples during the reprogramming process. **C**, Heatmap of the expression levels of of 562 genes associated with a consensus pluripotency signature (Chung et al., 2012). *miR-203*(+/+) (WT), *miR-203*(−/−) (KO) and *Cola1*(miR203)(KI). **D**, Expression of the indicated transcripts. Colors indicate expression level as shown in the scale to the right. The distribution of the samples is as in the PCA plot in panel B. **E**, Immunodetection of the indicated proteins in lysates obtained at different time points during reprogramming. GAPDH was used as a loading control.

### miR-203 over-expression prevents in vivo reprogramming

We next tested the effects of miR-203 expression in reprograming in vivo by combining the miR-203 mutant alleles with an inducible model in which the four reprogramming factors (tet-inducible OSKM cassette; Figure 2A) can be induced upon administration of doxycycline in vivo (Abad, Mosteiro et al., 2013). In line with the observations in vitro, miR-203 overexpression prevented in vivo reprogramming, whereas it was accelerated and enhanced in miR-203-knockout mice as detected by the deleterious effects in survival as a consequence of cell reprogramming in vivo (Figure 2B). The histopathological abnormalities observed in wild-type mice were very similar to those reported previously (Abad et al., 2013). Lack of miR-203 resulted in exacerbated alterations, including more pronounced dedifferentiation in the intestine, the stomach and other tissues, disorganization in different organs such as the kidney and liver, tumoral masses in kidney, liver and pancreas and liver anisocytosis (Figure 2C, D and Figure EV2). None of those abnormalities were observed in miR-203 overexpressing mice, which survived without obvious histological alterations (Figure 2B, C), indicating that this microRNA was able to prevent cellular and tissue reprogramming not only in vitro but also in vivo.

**Figure 2.**
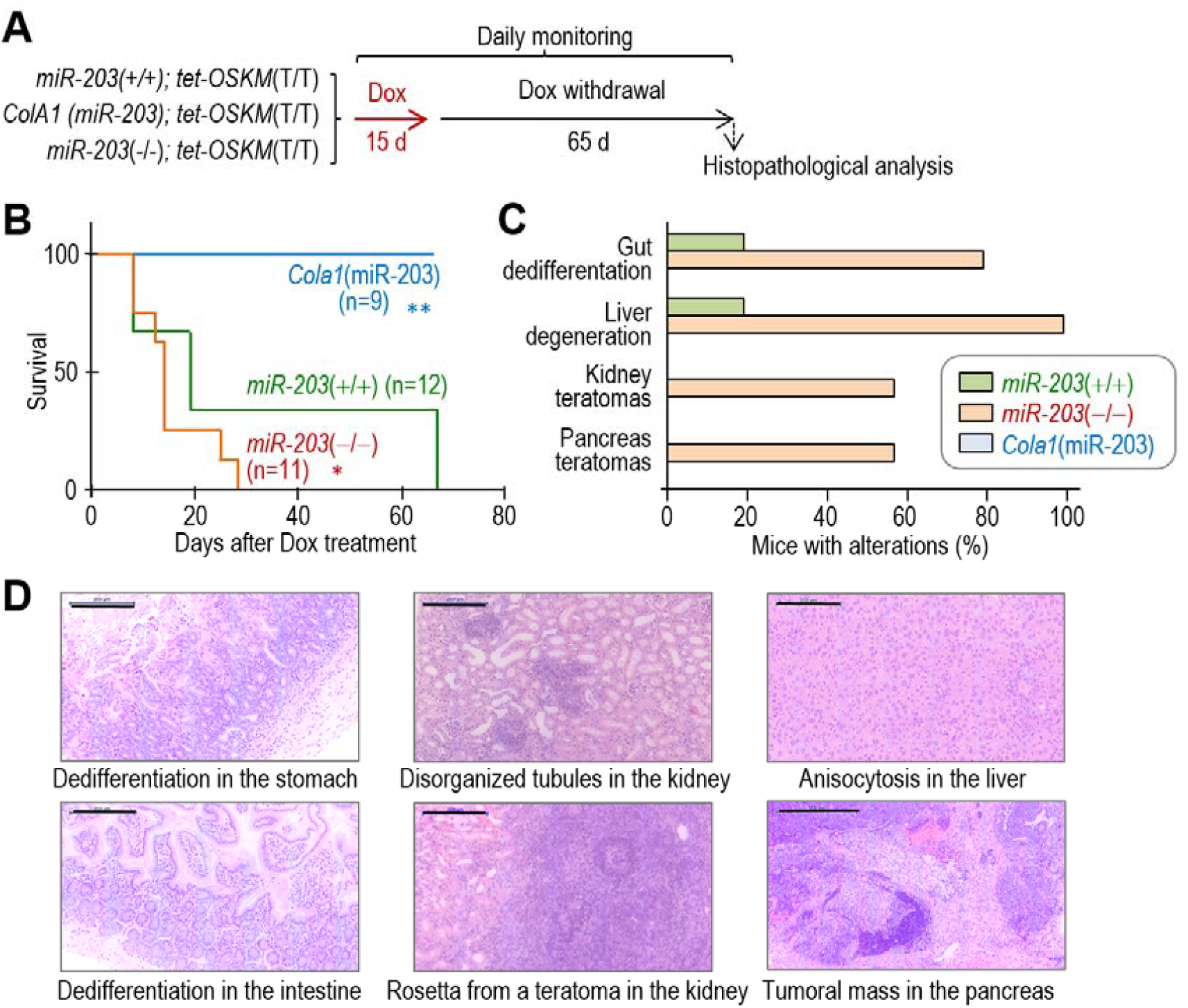
miR-203 prevents reprogramming in vivo. **A**, Schematic representation of the protocol followed for in vivo reprogamming in mice with the indicated genotypes. **B**, Survival curve of the indicated mouse strains in the presence of the tet-OSKM allele after doxycycline treatment during 15 days followed by Dox withdrawal. *, *P*<0.05; **, *P*<0.01 (One-way ANOVA test). **C**, Quantification of representative pathologies in the cohorts represented in panel A. See Figure S2 for details. **D**, Representative hematoxylin & eosin images of the most dramatic phenotypes in tet-OSKM; *miR-203*(−/−) mice after administration of Dox. Scale bars, 200 μm (500 μm bottom, right).

### miR-203 directly regulates *Nfatc2* expression during reprogramming

To analyze the mechanism by which miR-203 blocks reprogramming, we analyzed the transcriptomic profile of cells after exposure to the reprogramming factors in vitro (Figure 3A). We first compared the transcripts downregulated in *Cola1*(miR-203) iPSCs versus transcripts upregulated in *miR-203*(−/−) cells (Figure 3B). This resulted in a list of 29 transcripts, that revealed enrichment in a set of targets of the transcription factor NFATC2 (nuclear factor of activated T cells 2). We next searched for predicted miR-203 targets among the transcripts down-regulated in *Cola1*(miR-203) and up-regulated in *miR-203*(−/−) cells at different time points during the reprogramming process (Figure 3C). No transcript was common in the three independent combined lists generated at 4, 6 and 7 days, although *Dnmt3a* and *Dnmt3b*, two miR-203 targets previously validated in pluripotent cells (Salazar-Roa et al., 2019), were detected at day 7. Interestingly *Nfatc2* transcripts also present as predicted miR-203 targets significantly deregulated at day 6 and 7 (Figure 3C). In addition, transcription factor binding profile databases revealed NFATC2 as the only common hit regulating the genes significantly altered in all the comparisons tested (Appendix Table S3).

**Figure 3.**
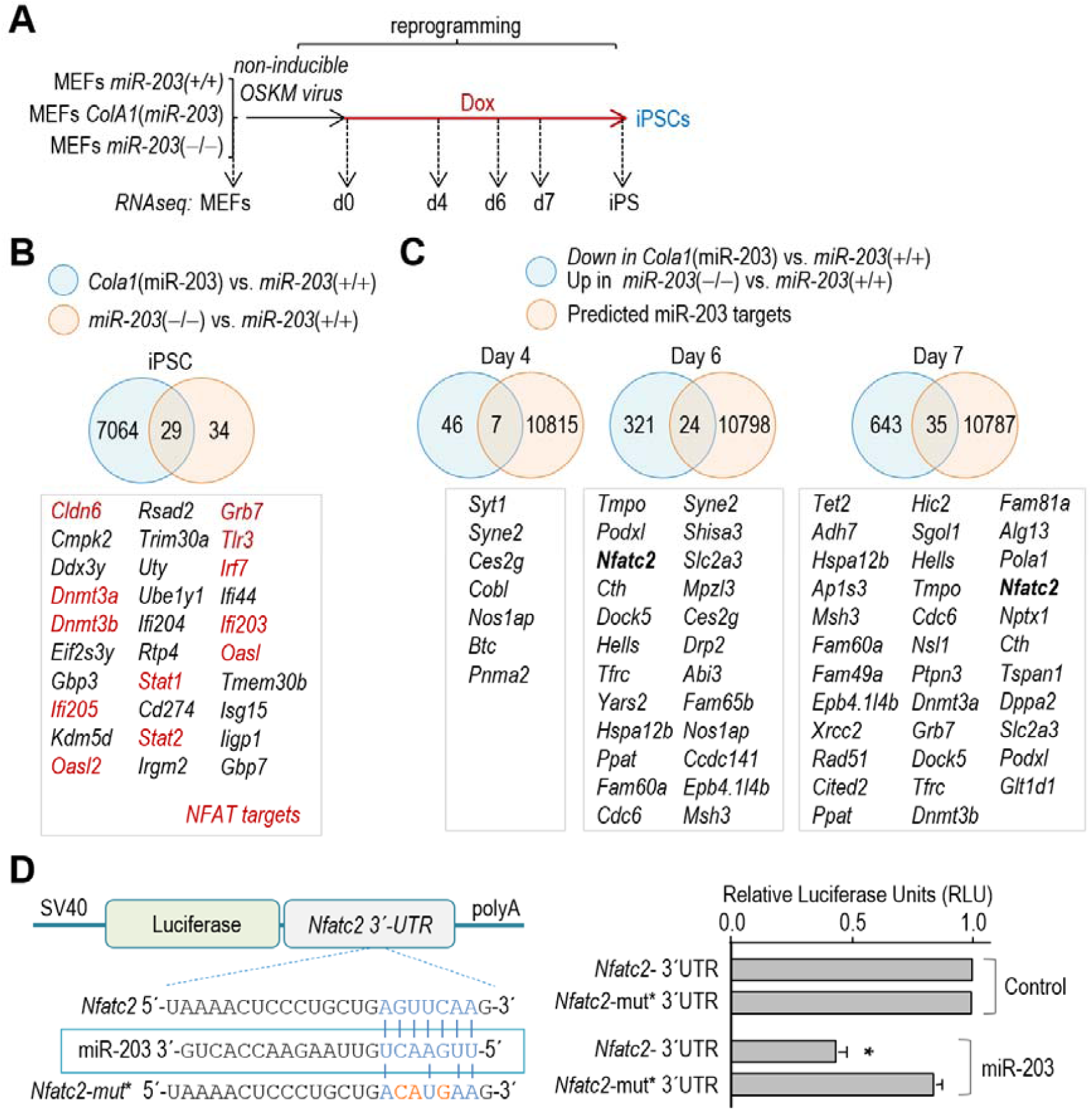
*Nfatc2* is a direct target of miR-203 deregulated during reprogramming. **A**, Schematic representation of the protocol followed for reprogramming and RNA analysis in vitro. **B**, Venn Diagrams showing genes significantly altered in *Cola1*(miR-203) (blue), and miR-203(−/−) (light brown) iPS cells. Potential NFATC2 targets are shown in red. **C**, Venn Diagrams showing the common genes significantly altered in the indicated comparisons: in blue, common genes down-regulated in *Cola1*(miR-203) and up-regulated in miR-203(−/−) cells; in light brown, predicted miR-203 targets. The list of common transcripts is shown below. **D**, The miR-203 target site contained in *Nfatc2* 3’-UTR is highlighted in blue and aligned with the corresponding miR-203 seed sequence. Mutated residues are indicated in red. The histogram shows Relative Luciferase Units (normalized to Renilla luciferase and relative to DNA amount) in 293T cells transfected with DNA constructs carrying the wild-type 3’UTR from *Nfatc2* or its mutated version as indicated, downstream of the luciferase reporter. Cells were co-transfected with *Renilla* luciferase as a control of transfection, together with miR-203 mimics or miRNA control. Data are represented as mean ± s.d. (n=2 independent experiments). *, *P*<0.05; Student’s t-test; unpaired; two-tailed).

To validate NFATC2 as a direct miR-203 target, we generated luciferase constructs fused to the *Nfatc2* 3’-UTR. Exogenous expression of miR-203 by miRNA mimics led to decreased luciferase signal and this effect was abrogated when the putative miR-203 binding sites at Nfatc2 3’-UTR were mutated, indicating a direct control of these transcripts by miR-203 (Figure 3D).

### NFATC2 is a critical target of miR-203 effects during reprogramming

In line with the reported biphasic role of NFATC2 on cellular preprogramming (Khodeer & Era, 2017, Sun, Liao et al., 2016), *Nfatc2* transcripts were slightly induced during reprogramming in wild-type cells (Figure 4A). These transcripts were repressed in *Cola1*(miR-203) cells and dramatically upregulated in *miR-203*(−/−) cells at intermediate stages during reprogramming to be finally silenced in stable iPS clones (Figure 4A). At the protein level, NFATC2 was induced by day 5 in wild-type cells as reported previously (Khodeer & Era, 2017, Sun et al., 2016), and this induction was enhanced in miR-203 knock-out cells (Figure 4B). Members of the NFAT transcription factor family, including NFATC2, are normally present in the cytoplasm in a hyperphosphorylated and inactive form. Increase in intracellular Ca2+ activate calcineurin, which dephosphorylates NFATs, promoting their activation and translocation to the nucleus, where they regulate NFAT-dependent transcription (Li, Rao et al., 2011). Subcellular fractionation of *miR-203*(−/−) cells confirmed the dephosphorylation and translocation of NFATC2 from the cytosol to the nucleus during reprograming (Figure 4C).

**Figure 4.**
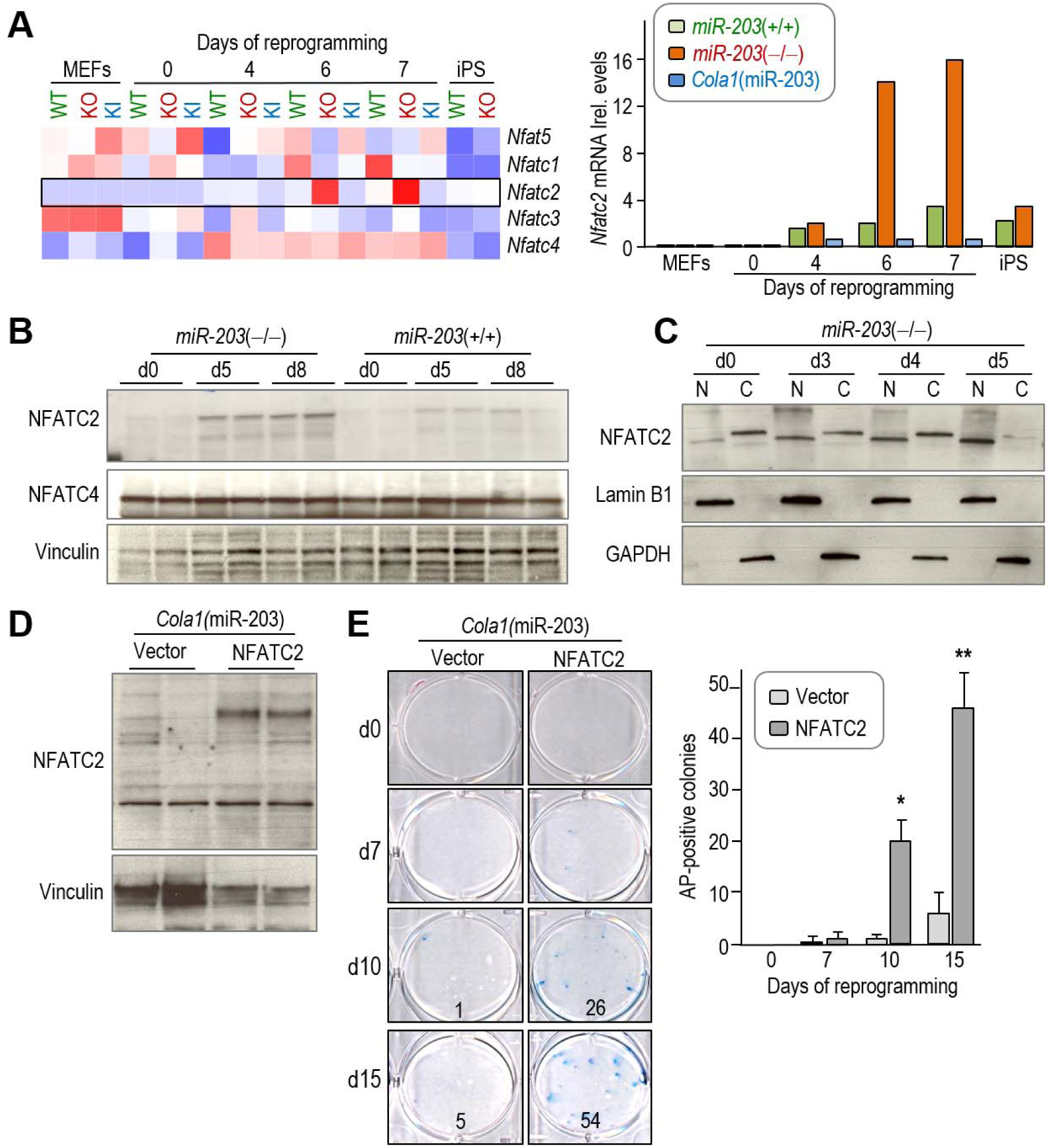
miR-203 regulates NFATC2 levels during cellular reprogramming. **A**, Heatmap plot showing the expression levels assessed by RNAseq of the NFAT family members (*Nfatc1, Nfactc2, Nfatc3, Nfatc4, Nfatc5*) during reprogramming. The histogram shows the validation of *Nfatc2* transcript levels by qRT-PCR. mRNA expression levels were normalized by the expression of *Gapdh* as housekeeping gene. Two independent experiments from two different MEF isolate lines are represented in the graph. **B**, Detection of NFATC2 and NFATC4 protein levels at different time points during reprogramming. Vinculin was used as a loading control. **C**, Subcellular fractionation analysis of NFATC2 in miR-203 knock-out cells. Lamin B1 is included as loading control for the nuclear fraction while GAPDH is included as loading control of the cytosolic fraction. For **B-C**, representative images of at least 3 independent experiments are shown. **D**, Detection of over-expressed transduced NFATC2 as determined by western blot, in miR-203 knock-in MEFs at day 7 of reprogramming. Vinculin protein levels are included as loading control. **E**, Left, Effect NFATC2 over-expression on cellular reprogramming of *Cola1*(miR-203). Representative images of 3 independent MEF isolates are shown. The total number of counted colonies at this particular representative experiment are indicated in the bottom right corner of each image. The graph to the right shows the quantification of the number of colonies as assessed by AP staining in the different samples. Values correspond to the average and s.d. (n= 3 independent MEF isolates per line). *, *P*<0.05; **, *P*<0.01; ***, *P*<0.001 (Student’s t-test; unpaired; two-tailed).

To investigate the functional relevance of NFATC2 during reprogramming, we first transduced *Cola1*(miR-203) cells with NFATC2 expressing vectors (Figure 4D). Remarkably, the exogenous expression of NFATC2 was sufficient to rescue the ability of *Cola1*(miR-203) cells to reprogram to iPSCs, as observed by the increase in the number of AP positive colonies at day 10 and day 15 of the reprogramming process (Figure 4E).

Furthermore, treatment of cells with (CsA), a calcineurin inhibitor (Figure 5A), resulted in decreased number of alkaline phosphatase-positive colonies, both in wild-type and *miR-203*(−/−) cultures (Figure 5B). CsA forms a complex with cyclophilin A, and it is this complex that binds and inhibits calcineurin (Schreiber & Crabtree, 1992). However, this complex has calcineurin-independent off-target effects that may account for side effects reported for this drug (Kiani et al., 2000; Martinez-Martinez & Redondo, 2004). NFAT activation by calcineurin can also be prevented by overexpressing peptides that span the PIXIT and the LxVP docking sites present in the NFAT regulatory domain (Li et al., 2011). We therefore used an alternative approach for calcineurin inhibition (Figure 5A) based on the expression of the LxVP peptide, which inhibits calcineurin independently of cyclophilin A to avoid off-target effects of CsA (Rodriguez, Roy et al., 2009). As represented in Figure 5C, transduction of wild-type and miR-203 knock-out cells, prior the reprogramming process, with lentiviruses encoding LxVP, but not with the AxAA mutant version (LxVP mut), significantly reduced the efficiency of reprogramming.

**Figure 5.**
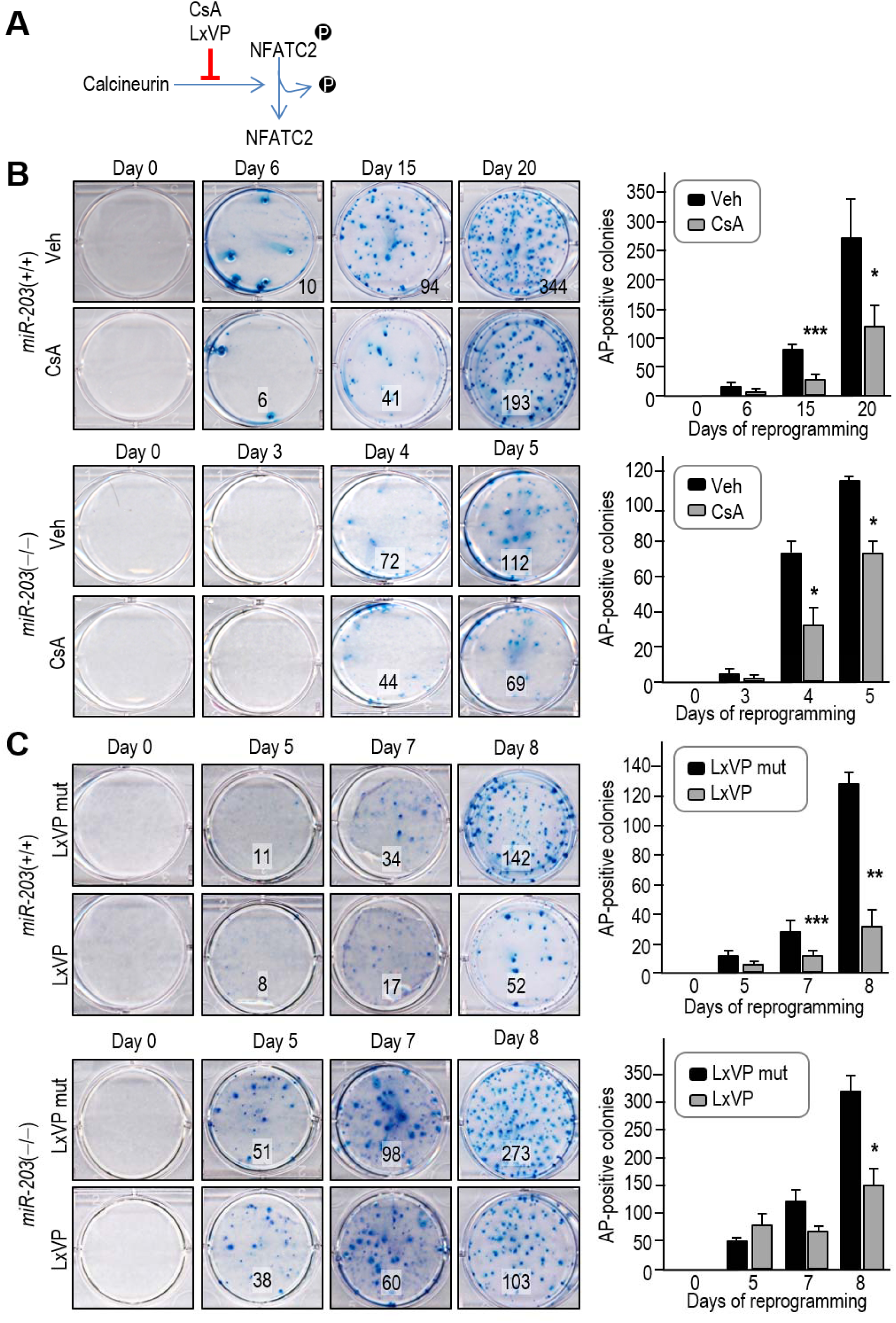
Inhibition of the calcineurin-NFATC2 pathway mimics the effect of miR-203 overexpression. **A**, Schematic representation of the calcineurin-NFATC2 pathway and the reagents used in this figure. **B**, Left, Effect of Cyclosporine A (CsA; 100ng/ml) during cellular reprogramming. Representative images of 3 independent MEF isolates per line are shown. Total number of counted colonies at this particular representative experiment are indicated in the bottom right corner of each image. The histogram to the right shows the quantification of the number of colonies as assessed by AP staining during the reprogramming process. Values correspond to the average and s.d. (n= 3 independent MEF isolates per line). *, *P*<0.05; **, *P*<0.01; ***, *P*<0.001 (Student’s t-test; unpaired; two-tailed). **C**, Effect of LxVP and LxVP mutant peptides on cellular reprogramming. Representative images of 3 independent MEF isolates per line are shown. Total number of counted colonies at this particular representative experiment are indicated in the bottom right corner of each image. The histogram to the right shows the quantification of the number of colonies as assessed by AP staining during the reprogramming process. Values correspond to the average and s.d. (n= 3 independent MEF isolates per line). *, *P*<0.05; **, *P*<0.01; ***, *P*<0.001 (Student’s t-test; unpaired; two-tailed).

Altogether, these results suggest that the miR-203-dependent repression of NFATC2 during reprogramming limits the ability of somatic cells to generate iPSCs in vitro and in vivo. An increasing number of evidences suggest that miRNAs are linked to pluripotency by controlling the expression of stemness transcription factors, epithelial-mesenchymal trans-differentiation, cell cycle progression or the epigenetic landscape of cells (Card, Hebbar et al., 2008, Leonardo, Schultheisz et al., 2012, Lin, Chang et al., 2008, Marson, Levine et al., 2008, Shenoy & Blelloch, 2014). Several microRNAs are associated to the acquisition of pluripotent features, such as those in the miR-302-367, miR-290-295 or miR-106-363 clusters (Card et al., 2008, Leonardo et al., 2012, Marson et al., 2008, Shenoy & Blelloch, 2014). miR-203, on the other hand, was originally found as a microRNA strongly accumulated in committed cells during skin differentiation (Lena et al., 2008, Wei, Orfanidis et al., 2010, Yi et al., 2008). Functional studies suggested that miR-203 repress the transcription factor p63 in addition to SKP2 and MSI2 thus limiting the stemness potential of skin progenitors (Jackson et al., 2013). Other studies in specific progenitor cell types suggest similar functions through the repression of additional targets such as GATA6, BMI1, ADAMDEC1 (Jimenez-Pascual, Hale et al., 2019, Wellner, Schubert et al., 2009).

NFAT family proteins are known to regulate lineage-specific transcription factors during differentiation of T helper 1 (Th1), Th2, Th17, regulatory T (Treg), and follicular helper T cells (Tfh) (Lee, Kim et al., 2018). Interestingly a biphasic role for calcineurin/NFAT signaling during reprogramming has been reported (Khodeer & Era, 2017, Sun et al., 2016). At early and decisive phases of reprogramming, calcineurin-NFAT signaling is transiently activated and such activation is required for effective reprogramming. Calcineurin is responsible for the dephosphorylation of multiple serine residues within the NFATC2 regulatory domain, thereby triggering nuclear translocation of NFATC2 to exert its function as transcription factor (Matsuda & Koyasu, 2000). Calcineurin-NFATC2 contributes to the reprogramming through regulating multiple early events, including the mesenchymal to epithelial transition (MET), cell adhesion and emergence of SSEA1 (+) intermediate cells, distinctive of the early steps on the reprogramming path. However, at later stages, calcineurin exerts a negative effect that is mediated by NFATC2. NFATC2 interacts with HDAC3, EZH2 and SUV39H1 to increase H3K9me3, and H3K27me3 over the *SOX2* enhancer and *KLF2* promoter, respectively, resulting in down-regulation of their expression. At this latest point, calcineurin/NFATC2 signaling becomes a barrier for pluripotency and NFATC2 is typically downregulated at later stages during reprogramming (Khodeer & Era, 2017, Sun et al., 2016).

Functional studies in miR-203-null or –overexpressing cells actually point to NFATC2 as a critical miR-203 target during pluripotency. First, miR-203 directly regulates *Nfatc2* transcripts through its 3’-UTR. Second, preventing the calcineurin-mediated dephosphorylation of NFATC2 with cyclosporin A (Handschumacher, Harding et al., 1984, Youn, Piao et al., 2002), or inhibiting the calcineurin-NFATC2 interaction mimic the effects of miR-203 overexpression. Finally, overexpression of NFATC2 rescues the defects of miR-203 overexpressing cells allowing the generation of iPS cultures. These data suggest NFATC2 as a major player in miR-203 effects but, given the promiscuous activities of microRNAs, we cannot rule out other possible miR-203 targets participating in this process. In fact, we found *Dnmt3a* and *Dnmt3b*, characterized recently as miR-203 targets (Salazar-Roa et al., 2019), as additional transcripts deregulated during reprogramming in a miR-203-dependent manner.

Despite the multiple observations suggesting miR-203 as a general repressor of the stemness potential, we recently found that transient expression of miR-203 in already-formed pluripotent cells enhance the differentiation and developmental potential of these cells towards multiple lineages (Salazar-Roa et al., 2019). These effects in pluripotent cells are due to widespread hypomethylation of the genome through the inhibition of the *de novo* DNA methylases *Dnmt3a* and *Dnmt3b*. This is fundamentally different of the effect of miR-203 during reprogramming, which seems to affect the early events in which calcineurin-NFATC2 are involved. Our data suggest that the biphasic dynamics of NFATC2 activity is limited by miR-203 during the early phases of reprograming, and eliminating miR-203 greatly improves reprogramming from fibroblasts in vitro. We speculate that this function may apply to many other cell types as elimination of miR-203 in adult mice dramatically enhances the effect of an inducible OSKM cassette during reprogramming in vivo. Altogether, these data suggest that transient miR-203 inhibition, e.g. using anti-miRNA sequences of microRNA sponges, may improve reprogramming in several cell types and could be incorporated to many protocols of use in regenerative medicine.

## Materials and Methods

### Animal models and procedures

Animal experimentation was performed according to protocols approved by the CNIO-ISCIII Ethics Committee for Research and Animal Welfare (CEIyBA). miR-203 knockout model was generated by flanking the mmu-mir203 locus with loxP using homologous recombination into ES cells (Figure EV1A). A frt-flanked neomycin-resistant cassette was used for selection of recombinant clones. Elimination of the neomycin-resistant cassette was achieved using a CAG-Flpe transgenic line (Rodriguez, Buchholz et al., 2000) and loxP sites were recombined using a EIIa-Cre transgene (Schwenk, Baron et al., 1995) resulting in the miR-203(−/−). The miR-203 inducible model has been described elsewhere (Salazar-Roa et al., 2019). These animals were maintained in a mixed C57BL6/J x 129 x CD1 genetic background. miR-203 mice were crossed with the i4F reprogrammable mice (Abad et al., 2013), including the doxycycline inducible tetracistronic cassette with the four murine reprogramming factors (Oct4, Sox2, Klf4, c-Myc). To induce the OSKM cassette in vivo, we followed the protocols described previously (Abad et al., 2013). Doxycycline (Sigma) was administered to mice in the drinking water supplemented with 7.5% of sucrose or alternatively orally in diet (Dox delayed release pellets, from Jackson laboratories) during 15 days, followed by Dox withdrawal during 65 more days. Mice were monitored daily and euthanized when any sign of suffering was noticed. For histological analsysis, tissue samples were fixed in 10% neutral buffered formalin (4% formaldehyde in solution), paraffin-embedded and cut at 3 µm, mounted in superfrost®plus slides and dried overnight. Consecutive sections were stained with hematoxylin and eosin (H&E). The images were acquired with a slide scanner (Axio Scan Z1, Zeiss). Images were captured and quantified using the Zen Software (Zeiss).

### Cell culture and gene expression

Primary mouse embryonic fibroblasts (MEFs) were obtained from embryos at E13.5 and cultured in DMEM supplemented with 10% of FBS and Penicillin/Streptomycin. Cultures were routinely tested for mycoplasma. Reprogramming in vitro was promoted on these MEF cultures by non-inducible Oct4-Sox2-Klf4-cMyc (OSKM) lentiviral transduction. For lentiviral transduction, we transfected HEK293T cells with FUW-OSKM (Addgene #20328) and packaging vectors using Lipofectamine 2000 (Invitrogen). Viral supernatants were collected twice a day on two consecutive days starting 24 h after transfection and were used to infect MEFs, previously plated at a density of 250.000 cells per well in 6-well plates. Previous to infection, polybrene was added to the viral supernatants at a concentration of 2 μg/ml. Infected MEFs were then cultured in iPSC medium, containing KO-DMEM (Gibco), 2-Mercaptoethanol 50 mM (Invitrogen), non-essential aminoacids MEM NEAA (Invitrogen), Lif (Leukemia Inhibitor Factor, ESGRO, Millipore), Penicillin and Streptomycin (5000 μg/ml, Invitrogen) and 20% knock-out serum replacement (KSR, Invitrogen). Medium was changed every 24 h and plates were stained for alkaline phosphatase activity to assure the efficiency of reprogramming (AP detection kit, Sigma-Aldrich). Once colonies were picked, iPSCs were cultured in iPSCs media over mitomycin C (Roche)-inactivated feeder cells. For inducing transient miR-203 over-expression in the cultures, *Cola1*(miR-203/miR-203); *Rosa26*(rtTA/rtTA) cells were treated with Dox (1 μg/ml; Invitrogen) during the reprogramming process. In this inducible system, we always test that insert expression is uniquely dependent on Dox, and becomes absolutely undetectable after Dox withdrawal (Salazar-Roa et al., 2019). As a control of the treatment itself, Dox was also added and tested in wild-type and miR-203 knock-out MEFs.

For transfection with mimics, we used miRIDIAN microRNA human hsa-miR-203a-5p (C-302893-00)/has-miR-203a-3p (C-300562-03) mimics or mimic transfection control (scramble sequences purchased from GE Dharmacon) with Dy547 (CP-004500-01), all of them from GE Dharmacon. Transfection was performed using either Dharmafect transfection reagent (Dharmacon) or Lipofectamine RNAiMAX (Invitrogen) following the manufacturer’s instructions. Transfection efficiency was evaluated 24 h post-transfection by Dy547 fluorescence, and experiments were then performed as indicated in the figures.

Luciferase reported assays were performed in HEK293T cells. Briefly, 200,000 cells per well were seeded on 6-well plates and the day after, cells were transfected using Lipofectamine 2000 (Invitrogen), following the manufacturer’s instructions. The 3’UTR region from the murine gene Nfatc2 was amplified by PCR using cDNA clones (pCMV-Sport6-*Nfatc2*). PCR products were verified by sequencing and ligated into the pGL3-Control vector (Promega), downstream of the luciferase reporter gene. Mutations in the miR-203 binding sites were generated by site-directed mutagenesis and subsequently verified by sequencing. Transfections were performed with either miR-203 or miRNA control mimics as described above, in combination with the pGL3-derived vectors, and Renilla as a control. Luciferase measurement was achieved 48 h post-transfection using a luminescence microplate reader (Biotek).

For NFATC2 over-expression experiments, full-length cDNA of NFATC2 was subcloned into the retroviral vector pMCSV-PIG (Abad et al., 2013) by restriction-directed subcloning, using pEFBOS_HA_NFATC2 (Rodriguez, Martinez-Martinez et al., 2005) as template. For retroviral transduction, we transfected HEK293T cells with the respective pMCSV-PIG vector expressing a GFP reporter (including either only GFP or NFATc2-GFP) and the packaging vector pCL-ECO, using Lipofectamine 2000 (Invitrogen). Viral supernatants were collected twice a day on two consecutive days starting 24 h after transfection and were used to transduce MEF cells. Preceding the infection, polybrene was added to the viral supernatants at a concentration of 2 μg/ml. For LxVP over-expression, we transduced MEF cells with the lentiviral vectors for LxVPwt/ LxVP mut described previously (Rodriguez et al., 2009) and lentiviral transduction as performed as indicated above. Finally, Cyclosporin A (Sigma-Aldrich) was used at 100 ng/ml, as indicated in the Figure.

### Western Blot

For western blotting, cells were lysed in RIPA or a buffer containing 50 mM Tris HCl, pH 7.5, 1 mM phenylmethylsulphonyl fluoride, 50 mM NaF, 5 mM sodium pyrophosphate, 1 mM sodium orthovanadate, 0.1% Triton X-100, 1 μg/ml leupeptin, 1 mM EDTA, 1 mM EGTA and 10 mM sodium glycerophosphate. Thirty micrograms of total protein were separated by SDS-PAGE and probed with primary antibodies against NFATC2 (Santa Cruz Biotechnology, sc-7296), NFATC4 (Cell Signaling, 2188), Vinculin (Santa Cruz Biotechnology, sc-25336), Lamin B1 (Santa Cruz Biotechnology, sc-374015) or GAPDH (Santa Cruz Biotechnology, sc-365062).

### Analysis of mRNA and microRNA levels

RNA/microRNA was extracted from cell or tissue samples with Trizol (Invitrogen) or by using the miRVana isolation kit (Thermo Fisher), following the manufacturer’s recommendations. Retrotranscription into cDNA was performed using M-MLV Reverse Transcriptase (Promega) following the manufacturer’s protocol. Quantitative real time PCR was performed using Syber Green Power PCR Master Mix (Applied Biosystems) in an ABI PRISM 7700 Thermocycler (Applied Biosystems). The housekeeping gene *Gapdh* was used for normalization. The oligonucleotide primers used in this manuscript for quantitative and semi-quantitative PCR are included in Appendix Table S4.

For RNAseq, total RNA was extracted using the miRVana miRNA isolation kit (ThermoFisher), following the manufacturer’s recommendations. Between 0.8 and 1 µg of total RNA was extracted from cells during reprogramming, with RIN (RNA integrity number) numbers in the range of 9 to 10 (Agilent 2100 Bioanalyzer). Poly A+ fractions were purified and randomly fragmented, converted to double stranded cDNA and processed using the Illumina’s “TruSeq Stranded mRNA Sample Preparation Part # 15031047 Rev. D” kit. The adapter-ligated library was completed by PCR with Illumina PE primers (8-11 cycles) and the resulting directional cDNA libraries were sequenced for 50 bases in a single-read format (Illumina HiSeq2000) and analyzed with nextpresso (Graña, Rubio-Camarillo et al., 2018). Reads were quality-checked with FastQC (http://www.bioinformatics.babraham.ac.uk/projects/fastqc) and aligned to the mouse genome (GRCm38/mm10) with TopHat-2.0.10 (Trapnell, Roberts et al., 2012), using Bowtie 1.0.0 (Langmead, Trapnell et al., 2009) and Samtools 0.1.19 (Li & Durbin, 2009), allowing two mismatches and five multihits. Transcripts assembly, estimation of their abundances and differential expression were calculated with Cufflinks 2.2.1 (Trapnell et al., 2012), using the mouse genome annotation data set GRCm38/mm10 from the UCSC Genome Browser (Rosenbloom, Armstrong et al., 2015). A false discovery rate (FDR) of 0.05 was used as threshold for significance in differential expression. Expression plots and heatmaps were built using Scanpy (Wolf, Angerer et al., 2018) and GENE-E (http://www.broadinstitute.org/cancer/software/GENE-E/index.html). The analysis of transcription factor targets was performed using JASPAR (Fornes, Castro-Mondragon et al., 2020). miRanda (www.microRNA.org) and TargetScan (www.targetscan.org) were used for the analysis of predicted miR-203 targets.

### Statistics

Samples (cells or mice) were allocated to their experimental groups according to their pre-determined type (cell type or mouse genotype) and therefore there was no randomization. Investigators were not blinded to the experimental groups (cell types or mouse genotypes). Normal distribution and equal variance was confirmed in the large majority of data and therefore, we assumed normality and equal variance for all samples. Based on this, we used the Student’s t-test (two-tailed, unpaired) to estimate statistical significance. For contingency tables, we used the Fisher’s exact test. Statistical analysis was performed using Prism (GraphPad Software, La Jolla, CA) or Microsoft Office Excel.

### Data availability

RNAseq data has been deposited in the GEO repository under accession number GEO GSE86653.

## Acknowledgements

We thank María Jose Bueno and Marta Gómez de Cedrón help with the generation of the mouse models, and Maria Abad and Manuel Serrano for reagents and helpful discussions. We are indebted to the members of the CNIO Cell Division and Cancer laboratory for their constant support and advice. M.S.R. was supported by the Asociación Española contra el Cáncer (AECC; 2012 AIOA120833SALA and 2018 INVES18005SALA) and a Juan de la Cierva contract from the Spanish Ministerio de Ciencia, Innovación y Universidades (MICIU). JMR lab is supported by the MICIU/AEI/FEDER, UE (RTI2018-099246-B-I00) and Comunidad de Madrid through the European Social Fund (ESF)-financed program AORTASANA-CM (B2017/BMD-3676). MM lab is supported by grants from AEI-MICIU/FEDER (RTI2018-095582-B-I00, SAF2017-92729-EXP and RED2018-102723-T), and the Comunidad de Madrid (B2017/BMD-3884). CNIO is a Severo Ochoa Center of Excellence (MICIU award SEV-2015-0510).

## Author Contributions

M.S.R. performed most of the in vitro and in vivo assays, with the help of M.A.F and M.T. in reprogramming assays and validation of *Nfatc2* as a target. S.M. and J.M.R. helped in the molecular characterization of NFATC2. O.G.-C. helped with the RNAseq analysis. M.M. conceived the project and M.S.R. and M.M. supervised the work, analyzed the data and wrote the manuscript with the help of all the authors.

## Figures

**Figure EV1.**
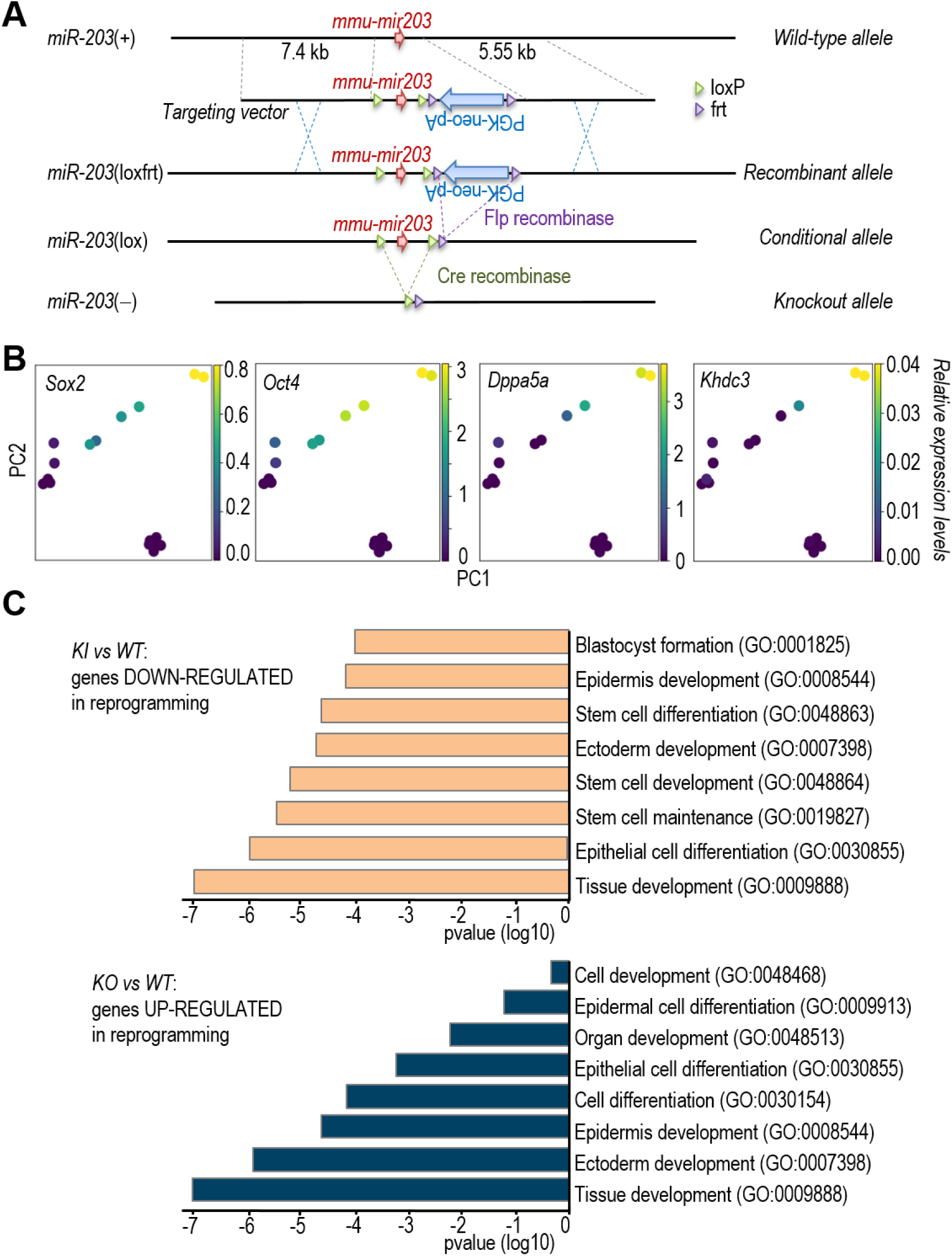
Mouse alleles generated in this work and effect of miR-203 on reprogramming. **A**, Schematic representation of the conditional knockout alleles generated at the mouse *mu-miR203* locus in this work. **B**, PCA plot of the indicated transcripts in the samples as represented in Figure 1. **C**, Top: Top categories in the Gene Ontology Analysis of the genes significantly down-regulated in miR-203-overexpressing cells when compared to wild-type cultures at day 7 of reprogramming (see also Table S1). Bottom: categories in the Gene Ontology Analysis of the genes significantly up-regulated in miR-203-null cells when compared to wild-type cultures at day 7 of reprogramming (see also Table S2).

**Figure EV2.**
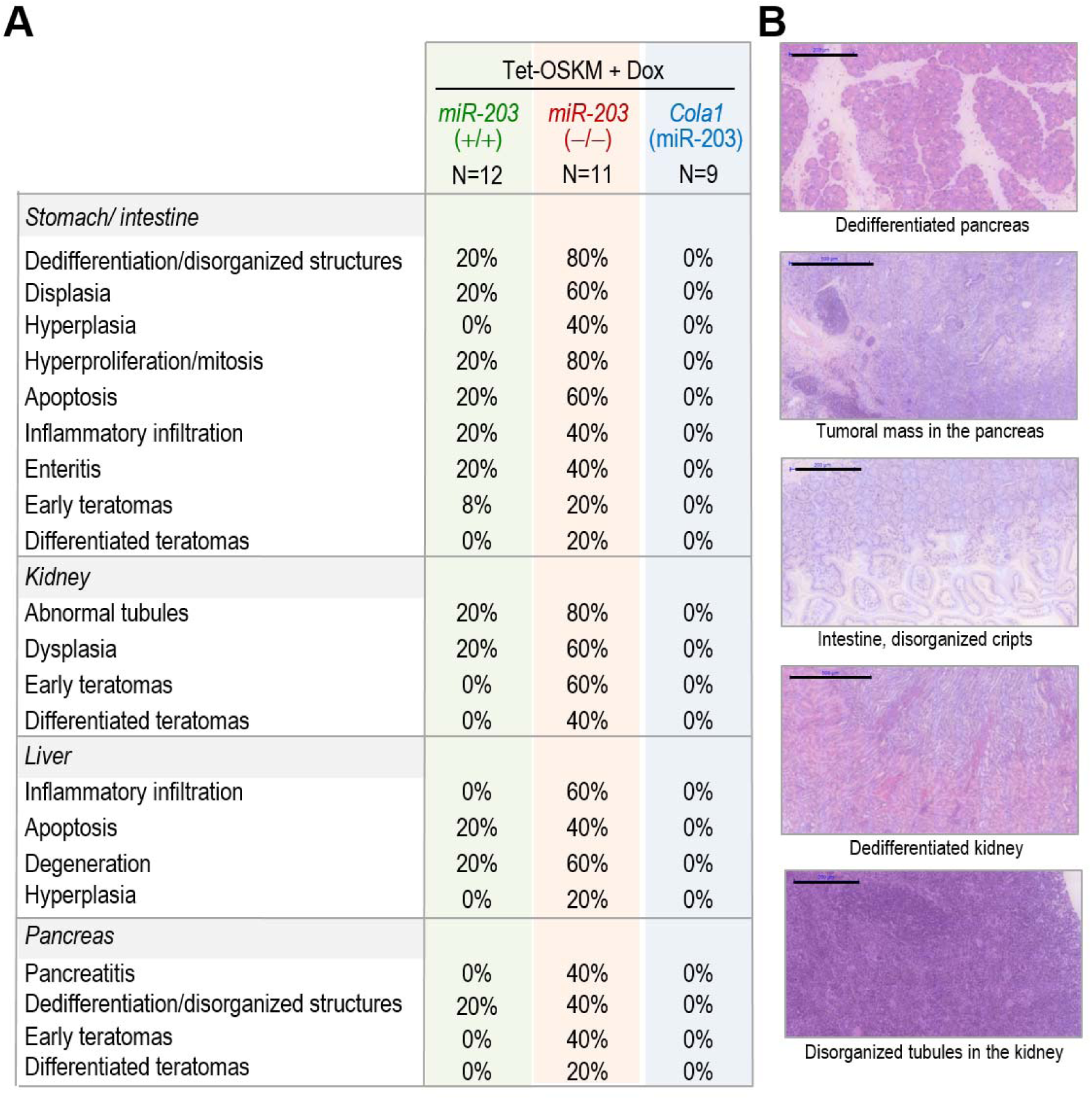
Pathologies observed during reprogramming in vivo in the indicated mouse models. **A**, Percentages of mice affected by the different pathologies indicated in the table. **B**, Hematoxylin & Eosin images of the most dramatic phenotypes observed in miR-203 knock-out mice. Scale bars, 200 μm (500 μm in the 2^nd^ and 4^th^ micrographs).

## Appendix Tables

**Appendix Table S1.** Gene ontology analysis of the genes significantly down-regulated in miR-203-overexpressing versus wild-type MEFs at day 7 of reprogramming to iPSCs.

| *Pathways* | *p-value*<br>*(log10)* | *Genes* |
| --- | --- | --- |
| tissue development<br>(GO:0009888) | -6.948 | *Gata4, Gjb3, Nanog, Erbb3, Pcdh8, Asprv1, Bmp8b, Upk1b, Gatm, Mylpf, Sprr1a, Krtdap, Tdgf1, Sfn, Krt14, Pou5f1, Krt17, Sall4, Ehf* |
| epithelial cell differentiation<br>(GO:0030855) | -5.804 | *Ezr, Ehf, Lama1, Sfn, Upk1b, Sprr1a, Sprr3, Krt14, Krt17, Elf3, Myb, Evpl* |
| stem cell maintenance<br>(GO:0019827) | -5.313 | *Nanog, Piwil2, Esrrb, Sox2, Pou5f1* |
| stem cell development<br>(GO:0048864) | -5.220 | *Nanog, Piwil2, Esrrb, Sox2, Pou5f1* |
| ectoderm development<br>(GO:0007398) | -4.750 | *Pou5f1, Lama5, Sfn, Krtdap, Sprr1a, Gjb3, Sprr3, Krt17, Ovol1, Evpl, Gli1* |
| stem cell differentiation<br>(GO:0048863) | -4.6683 | *Nanog, Piwil2, Esrrb, Sox2, Pou5f1* |
| epidermis development<br>(GO:0008544) | -4.220 | *Lama5, Sfn, Krtdap, Sprr1a, Gjb3, Sprr3, Krt17, Ovol1, Evpl, Gli1* |
| blastocyst formation<br>(GO:0001825) | -4.1772 | *Tead4, Esrrb, Pou5f1, Cdh1* |

**Appendix Table S2.**
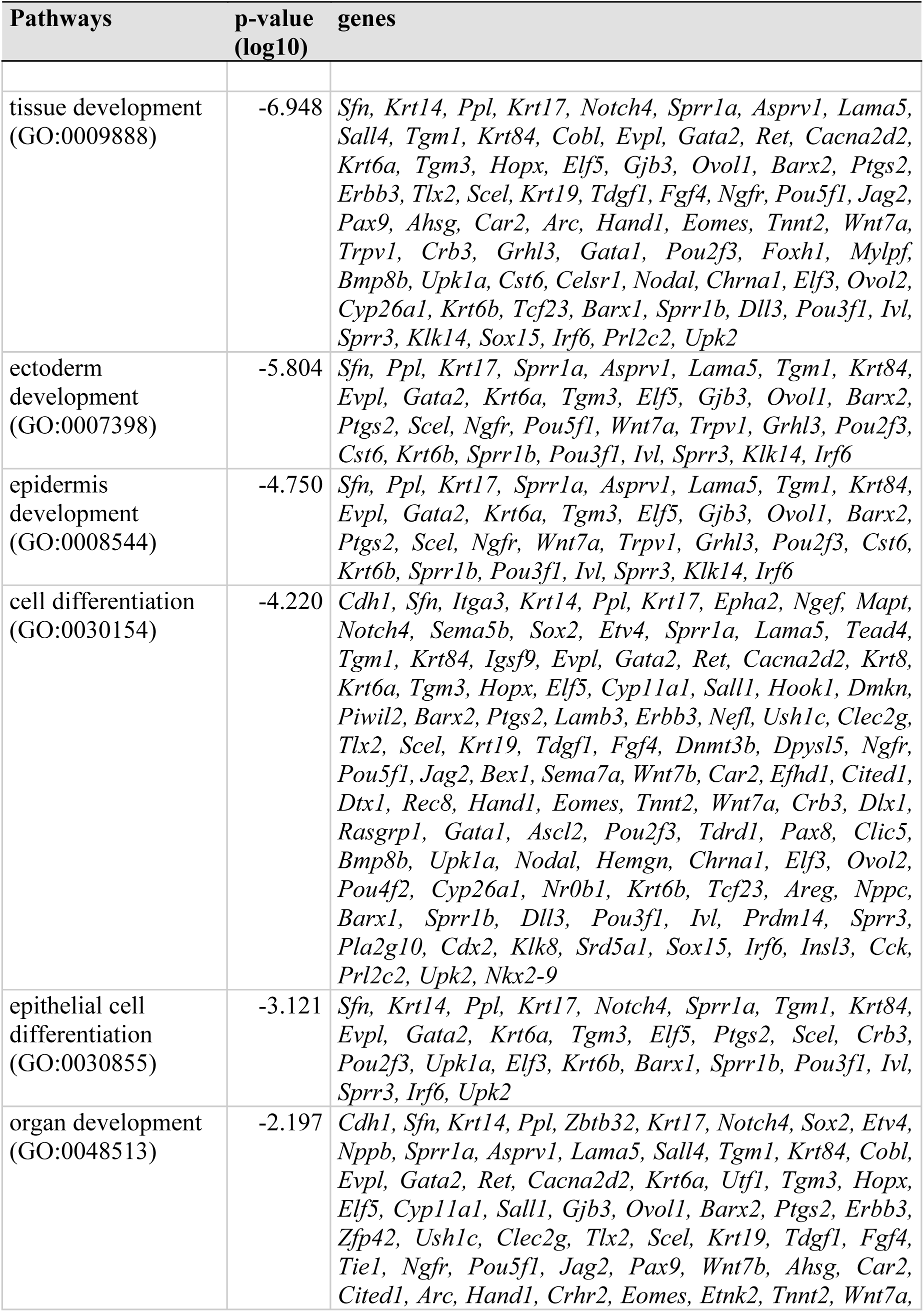
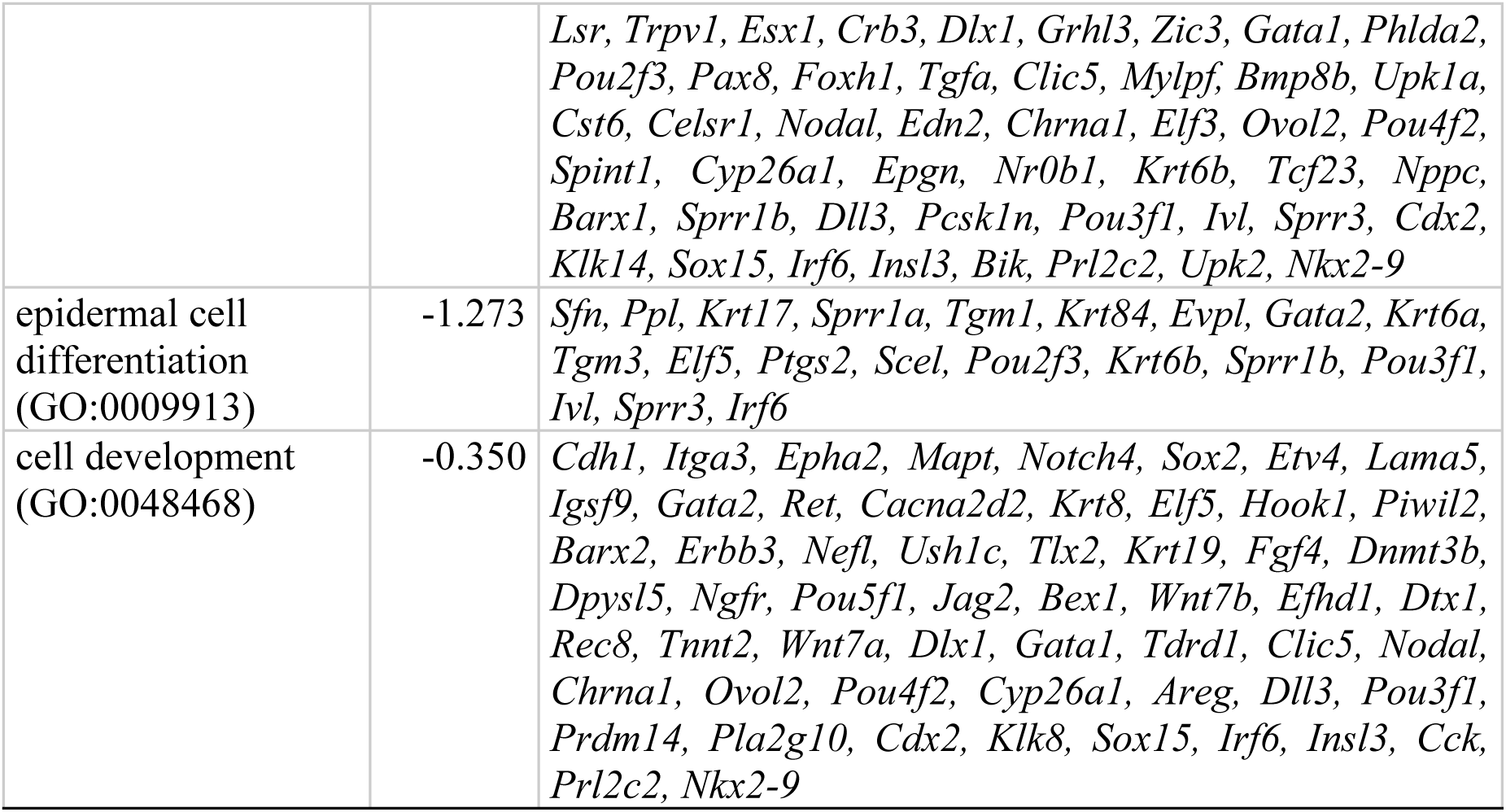
Gene ontology analysis of the genes significantly up-regulated in miR-203-null versus wild-type MEFs at day 7 of reprogramming to iPSCs.

**Appendix Table S3.**
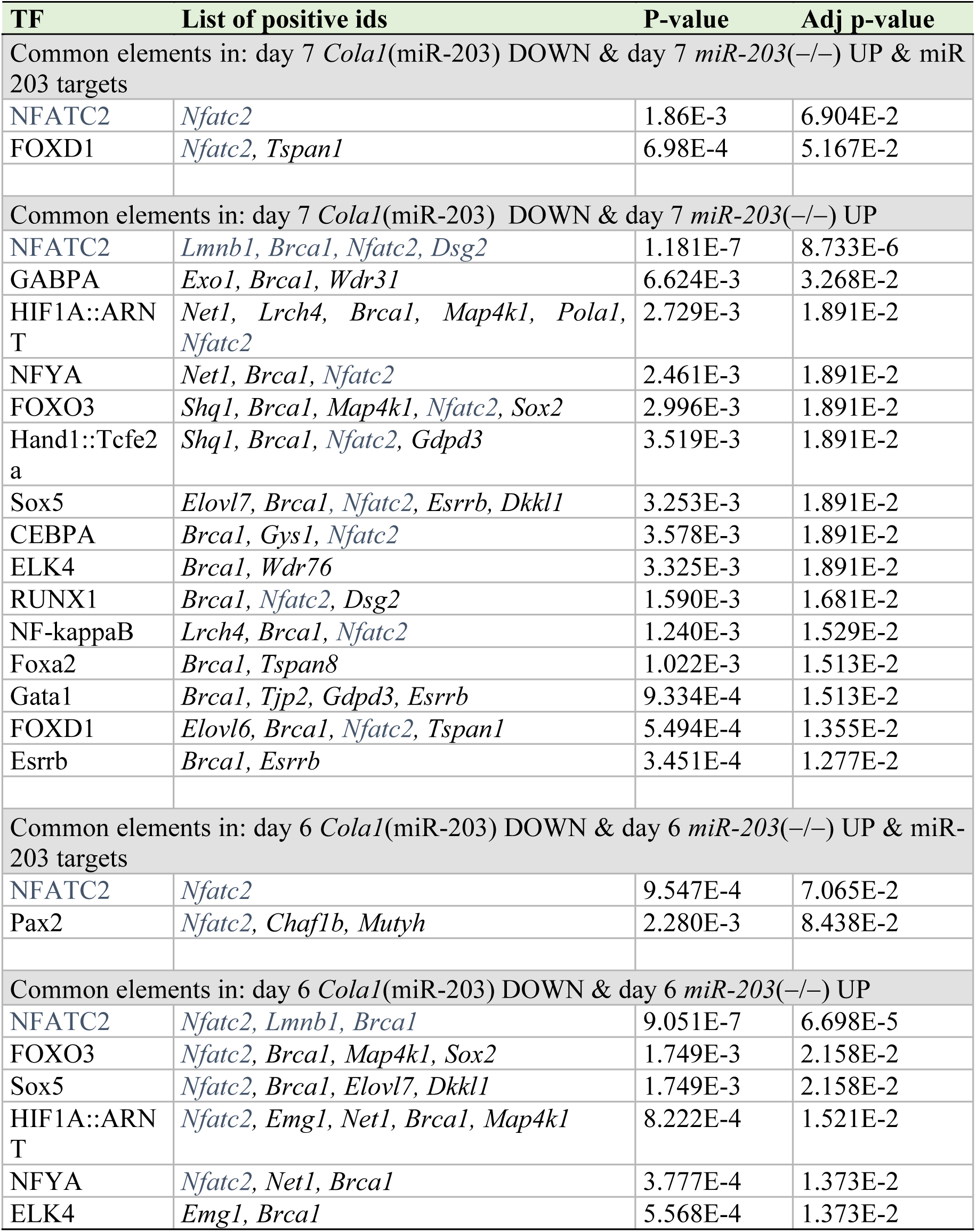
List of potential transcription factors regulating the genes significantly altered in the indicated comparisons (from Jaspar; transcription factor binding profile database). NFATC2 and NFATC2 targets are indicated in blue.

**Appendix Table S4.**
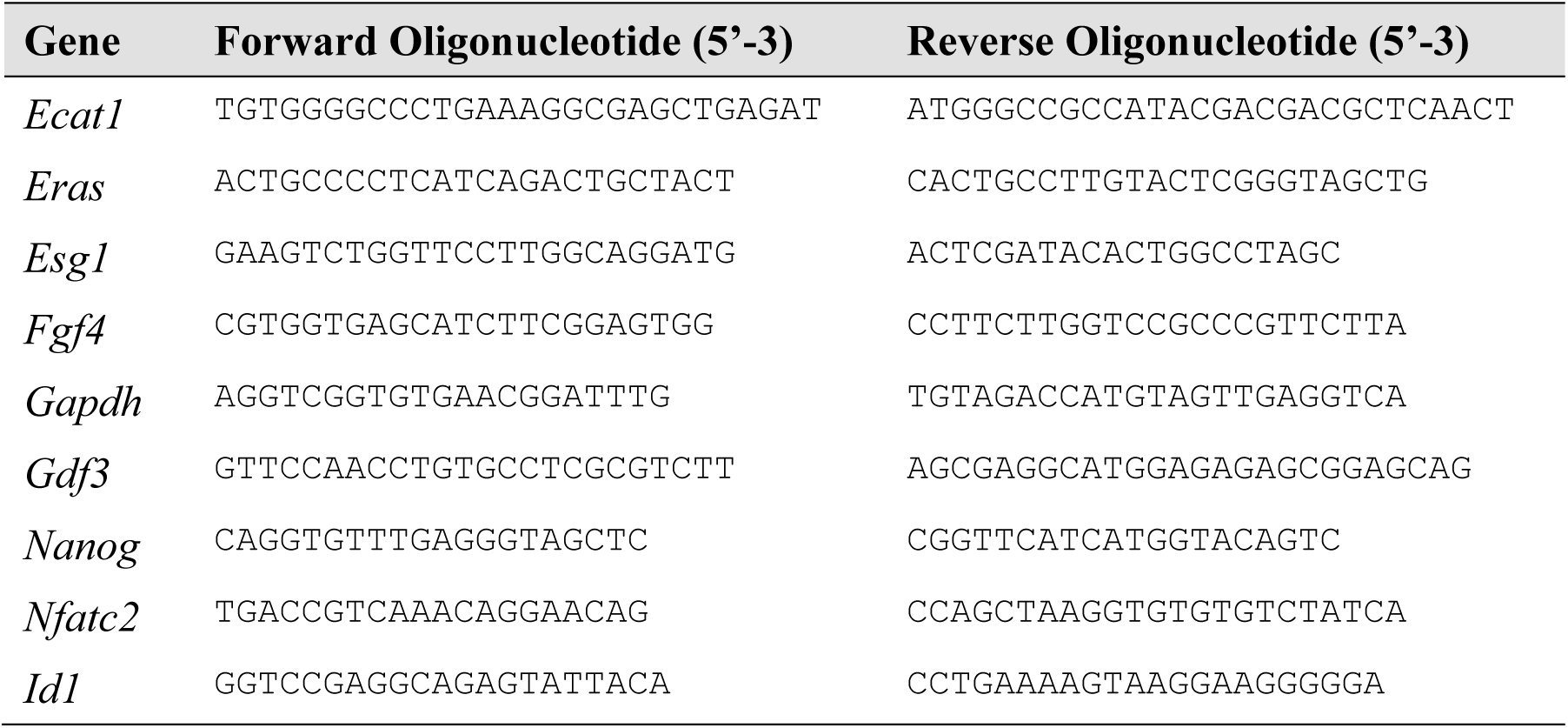
Oligonucleotides used in this work.

